# Simultaneous passive acoustic monitoring uncovers evidence of potentially overlooked temporal variation in an Amazonian bird community

**DOI:** 10.1101/2021.09.25.461815

**Authors:** W. Alexander Hopping, Christopher J. Sayers, Noe Roger Huaraca-Charca, Holger Klinck

**Affiliations:** K. Lisa Yang Center for Conservation Bioacoustics, Cornell Lab of Ornithology, Cornell University, Ithaca, NY, USA; Department of Ecology and Evolutionary Biology, University of California, Los Angeles, CA 90095, USA; Inkaterra Asociación, Victor Larco Herrera 130, Miraflores, Lima, Peru

**Keywords:** avian biodiversity, bioacoustics, passive acoustic monitoring, Amazon rainforest, temporal variation

## Abstract

The vocal activity and detectability of tropical birds are subject to high levels of temporal heterogeneity, but quantifying patterns of diel and day-to-day variation in complex systems is challenging with traditional point count methods. As a result, research concerning stochastic temporal effects on tropical avian assemblages is limited, typically offering only broad conclusions, e.g., overall activity is highest in the first few hours of the morning and some species are active at different times of the day. Passive acoustic monitoring introduces several advantages for studying temporal variation, particularly by enabling simultaneous and continuous data collection across adjacent sites. Here, we employed autonomous recording units to quantify temporal variation in avian vocal activity and observed species richness at an Amazonian reserve in Madre de Dios, Peru—a region featuring some of Earth’s richest, most complex avian assemblages. We manually annotated 18 dawn hour recordings, collected simultaneously from three separate days at the same six sites, which represent various microhabitats and avian community compositions. We documented significant and consistent temporal variation in avian vocal activity levels and observed species richness within the dawn hour and across days. We found that temporal effects were stronger for vocal activity than for observed species richness and that vocal activity patterns over the course of the dawn hour varied on a species-to-species basis. Our results indicate that overlooked temporal variation in Amazonian soundscapes may obfuscate the results of surveys that fail to sufficiently account for temporal variables with simultaneous monitoring. While manual analysis of large volumes of soundscape data remains challenging, such data should be collected as a supplement to traditional surveys whenever possible. Rapidly advancing research concerning the automated processing of acoustic data could lead to more efficient methods for reducing temporal bias and improving the calibration and accuracy of ornithological surveys in the tropics.

## Resumen

La actividad vocal y la detectabilidad de las aves tropicales están sujetas a altos niveles de heterogeneidad temporal, pero cuantificar con precisión los patrones de variación diurna y diaria en sistemas complejos es un reto con los métodos tradicionales de conteo por puntos. En consecuencia, la investigación sobre los efectos temporales estocásticos en los conjuntos de aves tropicales es limitada y normalmente sólo ofrece conclusiones generales, como que la actividad general es mayor en las primeras horas de la mañana y que algunas especies están activas en distintos momentos del día. El monitoreo acústico pasivo presenta varias ventajas sobre los métodos tradicionales para estudiar la variación temporal, en particular al permitir el recojo simultáneo de datos en lugares adyacentes. En este trabajo empleamos unidades de grabación autónomas para cuantificar la variación temporal en los niveles de actividad vocal de las aves y la riqueza de especies observada en una reserva amazónica en Madre de Dios, Perú, una región que cuenta con algunos de los conjuntos de aves más ricos y complejos del planeta. Anotamos manualmente 18 grabaciones de la hora del amanecer, recogidas simultáneamente en tres días distintos en los mismos seis lugares, que representan una variedad de microhábitats y composiciones de comunidades de aves. Se documentó una variación temporal significativa y consistente en los niveles de actividad vocal de las aves y en la riqueza de especies observada dentro de la hora del amanecer y a lo largo de los días. Descubrimos que los efectos temporales eran mayores en la actividad vocal que en la riqueza de especies observada, y que los patrones de actividad diurna variaban de especie a especie a escala fina. Nuestros resultados indican que la variación temporal que se pasa por alto en los paisajes sonoros amazónicos puede ofuscar los resultados de los estudios que no tienen suficientemente en cuenta las variables temporales con un seguimiento simultáneo. Nuevas investigaciones sobre el tratamiento automatizado de los datos acústicos podrían conducir a métodos más eficaces para reducir el sesgo temporal y mejorar la calibración y precisión de los estudios ornitológicos en los trópicos.

## 1 Introduction

The tropics account for an overwhelming share of Earth’s avian diversity, featuring more than 75% of all species and over 90% of terrestrial birds (Barlow *et al*. 2018). Still, tropical regions are underrepresented in ornithological literature relative to temperate regions, even though tropical birds have fundamentally different life history strategies, behavioral ecology, and vocal patterns than temperate birds, requiring separate study (Stutchbury & Morton 2008). The largest intact tropical system on the planet is the Amazon basin (Allan *et al*. 2017), which is the epicenter of global biodiversity (Antonelli *et al*. 2018). Like most of the tropics, the Amazon is undergoing rapid ecological change, primarily the result of agricultural expansion (Lapola *et al*. 2023), exacerbated by the increasing impacts of climate change (Xu *et al*. 2020). These anthropogenic impacts are significantly outpacing natural processes in Amazonia (Albert *et al*. 2023), threatening the resilience and stability of the region (Lovejoy & Nobre 2019, Boulton *et al*. 2022). Despite its importance, however, the Amazon’s avifauna remains poorly documented and is subject to flawed baseline species occurrence data and significant knowledge gaps (Lees *et al*. 2014), which are exacerbated by systemic barriers to researchers from the Global South (Soares *et al*. 2022). Efforts to increase ornithological survey coverage in the region are complicated by the difficulty of tropical field surveys. Overwhelming avian species richness, including many rare and similar species, challenging logistics, and poor visibility conditions contribute to the unreliability of avian surveys in Amazonia, where upwards of 95% of birds are heard but never seen by a field observer (Robinson *et al*. 2018).

The vocal activity and detectability of Neotropical birds are subject to high levels of temporal heterogeneity. Possible drivers of this variation include seasonality (Pérez-Granados & Schuchmann 2022), weather and feeding opportunities (Metcalf *et al*. 2021), foraging strata (Berg *et al*. 2006), temporal partitioning to avoid signal masking from other birds (Planqué & Slabbekoorn 2008, Luther 2008, Luther 2009, Hart *et al*. 2021) and vocalizing insects (Hart *et al*. 2015, Alvarez-Berríos *et al*. 2016, Aide *et al*. 2017, Metcalf *et al*. 2020), along with a wide range of poorly-understood floristic, geographic, and spatial variables (Menger *et al*. 2017). Some broad patterns of diel variation in the detectability and vocal activity of tropical birds are well established; for example, birds are most vocally active during the first 2–3 hours after dawn (Lynch 1995, Woltmann 2005). However, the temporal variation of tropical soundscapes is more complex than hour-long intervals can capture (Verner & Ritter 1986, Rodriguez *et al*. 2014, Metcalf *et al*. 2021). Many Amazonian species are only aurally detectable within strict temporal niches (Gil & Llusia 2020). These species may only vocalize during a single 5-minute period in the morning, perhaps accompanied by an even shorter bout in the evening (Parker 1991). Some Amazonian species have been shown to sing as infrequently as twice in 50 days (Jirinec *et al*. 2018). In addition to affecting estimates of total species richness, this fine-scale temporal variation in vocal activity can also influence patterns of observed community composition. Temporal effects on detectability vary by functional or taxonomic group. For example, canopy species are known to reach their activity peak later in the morning than understory species (Blake 1992), and species with high sensitivity to habitat fragmentation may have proportionately higher detection rates in pre-dawn surveys than less sensitive species (Woltmann 2005).

Quantifying the fine-scale patterns of temporal variation in tropical bird communities, both within and across days, is difficult. Non-simultaneous survey methods are subject to a litany of spatial, temporal, and observer biases, and therefore, capturing such granular effects can require infeasibly large sample sizes (Lynch 1995). For example, Esquivel & Peris (2008) found that four visits per point are necessary to account for the temporal variation in bird activity in the Atlantic Forest of Paraguay. This sampling design does not account for possible variation in vocal activity levels between different days, between different observers (Robinson *et al*. 2018), nor travel time between sites. Additionally, Rodriguez *et al*. (2014) found that temporal effects on acoustic activity patterns were spatially heterogeneous at adjacent recording sites in French Guiana. Therefore, precise assessments of temporal variation may be required on a per-site basis.

More costly and labor-intensive approaches, for example, using a team of point count technicians to visit multiple points simultaneously, may also introduce observer effects. Amazonian soundscapes include thousands of vocalization types, inducing high rates of false-negative and false-positive identification errors (Remsen 1994), and it can take months for skilled observers to reach a level of competence attainable in just one to two weeks of field experience in temperate systems (Parker 1991). Observer error rates have been experimentally quantified in relation to aural identification (Simons *et al*. 2007), visual identification (Hull *et al*. 2010), and the subjectivity of abundance estimates (Cerqueira *et al*. 2013). These dynamics are not limited to inexperienced surveyors; established ornithologists can also be subject to non-trivial identification error rates in environments as complex as Amazonia (Lees *et al*. 2014).Overlooked species tend to reflect non-random subsets of assemblages, and the specific nature of these biases may differ between individual observers, even amongst experts (Blackburn & Gaston 1998). Therefore, standardized surveys that do not produce archivable raw data and seek to uncover subtle spatiotemporal variation in avian vocal activity, observed species richness, and community structure should ideally be conducted by a single observer (Blake & Loiselle 2015, Robinson *et al*. 2018).

Simultaneous passive acoustic monitoring (PAM) conducted with autonomous recording units (ARUs) can eliminate much of the temporal bias in surveys by collecting continuous data from adjacent locations simultaneously (Venier *et al*. 2012, Tegeler *et al*. 2012). Such effects are difficult to quantify with traditional surveys, and establishing their presence and magnitude may be useful for improving the calibration of point count studies, even with relatively small volumes of ARU data. PAM can also address observer bias since data collection is independent of technician skill level, and recordings collected simultaneously can be annotated by a single observer, replayed and reanalyzed, archived publicly, and distributed to experts for an independent review of identifications (Rempel *et al*. 2005, Robinson *et al*. 2018). PAM has been shown to present a suitable supplement or alternative to point counts, both in Amazonia (Haselmayer & Quinn 2000), and in general (Shonfield & Bayne 2017, Darras *et al*. 2019, Blake 2021), while providing numerous other advantages (Acevedo & Villanueva-Rivera 2006, Newson *et al*. 2017, Darras *et al*. 2018, Jorge *et al*. 2018, Pillay *et al*. 2019, Sugai & Llusia 2019).

This study aimed to investigate the hypothesis that Amazonian soundscapes may be subject to overlooked temporal variation in avian vocal activity, both between days and throughout a single dawn hour, which in the absence of simultaneous survey methods, could mask important differences in vocal activity levels, community composition, and observed species richness between sites. We tested this hypothesis by deploying an array of ARUs at a single Amazonian reserve in Madre de Dios, Peru, and manually annotated a subsample of the resulting bioacoustic data. We predicted that stochastic effects of diel and day-to-day variation on vocal activity and observed species richness would be consistent across sites and, therefore, would be possible to control for with simultaneous monitoring.

## 2 Materials & Methods

### 2.1 Study area

We conducted this study at Inkaterra Reserva Amazónica (ITRA) in the Madre de Dios Department of southeastern Peru (12°32’07.8” S, 69°02’58.2” W). This 191-ha private reserve is located at an elevation of approximately 200 m along the Madre de Dios River, directly across from the Reserva Nacional de Tambopata—one of the most biodiverse regions on the planet (Foster *et al*. 1994) (Fig. 1). The habitat at ITRA primarily consists of *várzea* floodplain forest, the most endangered forest type in the southwestern Amazon (Phillips *et al*. 1994), interspersed with seasonally flooded *Mauritia* palm swamp forest.

**Figure 1:**
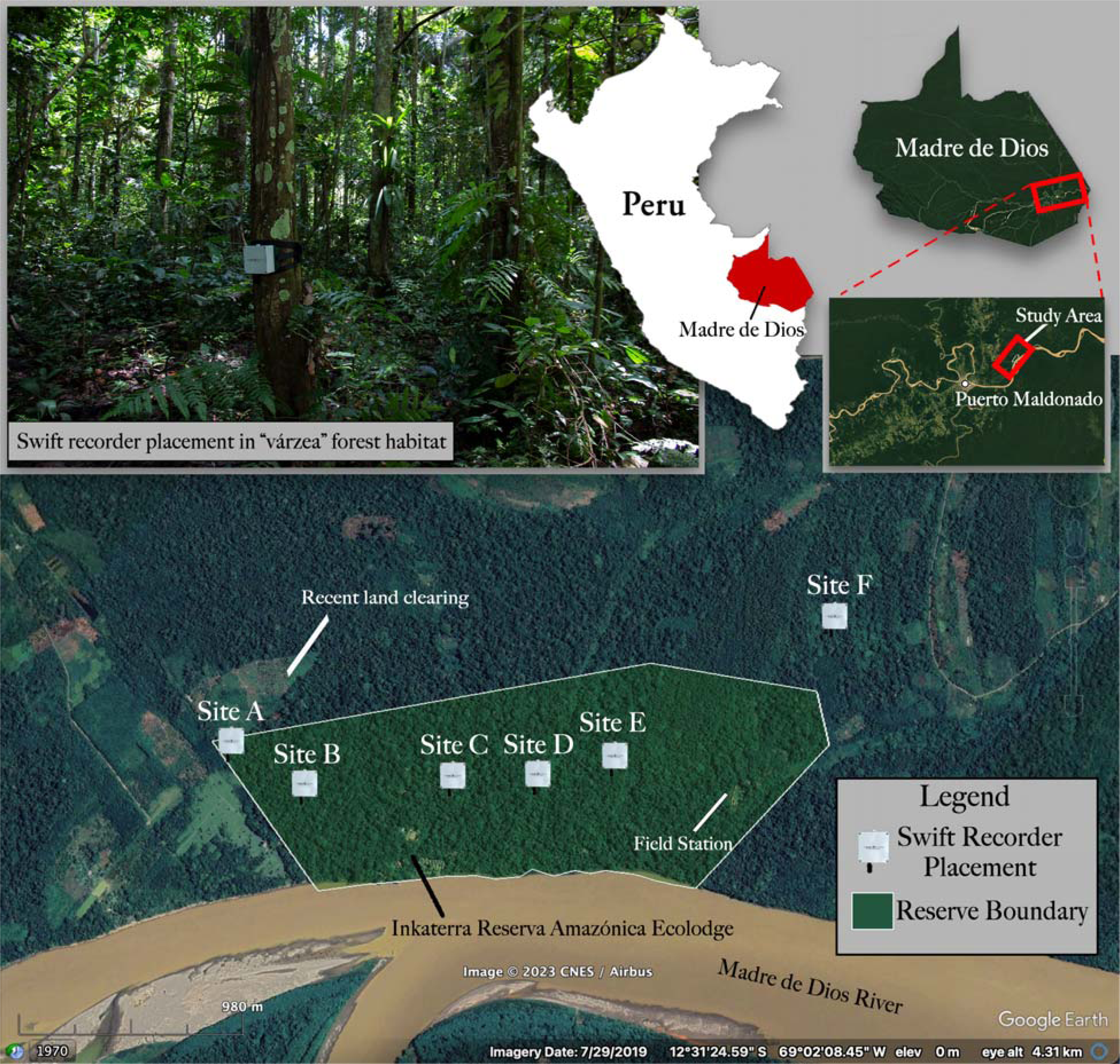
Swift recorder locations and typical forest habitat at Inkaterra Reserva Amazónica, Madre de Dios, Peru. Recorder locations covered a range of habitats, including edge habitat adjacent to clearings for small-scale agriculture (Site A), forest degraded by selective logging outside of the reserve’s boundaries (Site F), mature *várzea* in the reserve interior (Site C and Site E), and mixed palm swamp (*aguajal*)-*várzea* habitat (Site B and Site D).

### 2.2 Acoustic monitoring

We collected acoustic data from 14 January to 2 February 2019, using six ARUs (Swift recorder, K. Lisa Yang Center for Conservation Bioacoustics, Cornell Lab of Ornithology, Cornell University, Ithaca, NY, USA). We deployed the ARUs across the entire reserve, at a minimum distance of 350 m from each other and 450 m from the river, to limit spatially overlapping detections (pseudoreplication) and background noise (Ralph *et al*. 1995, Yip *et al*. 2017, Darras *et al*. 2018, Haupert *et al*. 2022), respectively. Deployment locations were chosen to represent a gradient of intactness and forest habitat type, including edge habitat adjacent to clearings for small-scale agriculture (Site A), forest degraded by selective logging outside of the reserve’s boundaries (Site F), mature *várzea* in the reserve’s interior (Site C and Site E), and mixed palm swamp (*aguajal*)-*várzea* habitat (Site B and Site D) (Fig. 1). Placed at the height of 1.5 m above the ground to maximize the sound detection space and minimize sound shadows (Darras *et al*. 2018), the 6 ARUs recorded continuously (mono, WAV format) throughout the deployment period using a sampling rate of 48 kHz (16-bit resolution) and a gain setting of 35 dB. The microphone sensitivity of the Swift was –44 dBV/Pa (+/– 3 dB) and featured a flat frequency response (+/– 3 dB) in the frequency range of the vocalizations of interest. The clipping level of the analog-to-digital (ADC) converter was +/– 0.9 V.

### 2.3 Annotation process

We manually annotated eighteen total dawn hours, from 05:00–06:00 PET (10:00–11:00 UTC), representing six sites on three days, using the Raven Pro Sound Analysis Software (version 1.6; K. Lisa Yang Center for Conservation Bioacoustics, Cornell Lab of Ornithology, Cornell University, Ithaca, NY, USA). Annotations were performed by a single observer, the lead author, to control for observer bias. A local expert, Noe Huaraca-Charca, also annotated a subset of these recordings (n = 4) as a quality control measure. The three days, 16 January (Day 1), 20 January (Day 2), and 31 January (Day 3), 2019, were randomly chosen from a subset of days that featured less than 5 min of heavy rain during the dawn hour. The dawn hour was selected because it typically contains the highest level of vocal activity and diversity among Neotropical birds (Berg *et al*. 2006), including many species that only sing in narrow time windows around first light (Parker 1991, Blake 1992). Additionally, the dawn soundscape in Amazonia features far less anthropogenic activity and insect noise than in other periods, reducing interference with avian species detection (Metcalf *et al*. 2020). While annotating short, randomly-selected samples spread out over a longer survey period can improve estimates of alpha and gamma diversity (Wimmer *et al*. 2013, Metcalf *et al*. 2021), this study was designed to assess temporal variation over the course of a contiguous hour, and annotations were therefore conducted in this format.

Spectrograms were viewed in 60 s increments split over two rows (Hann window, 3 dB bandwidth = 100 Hz, 690 FFT window size, 1024-sample DFT, 95% temporal overlap) (Fig. 2). To reduce bias introduced by improved sound identification knowledge developed throughout the project, the order in which recordings were annotated was determined using a random number generator. Vocalizations were labeled for both species and a handful of broader groups with confusing or indistinguishable calls (e.g., “TRSP” as “trogon species” [Trogonidae sp.]). Background vocalizations, audible but not clearly visible on a spectrogram, were denoted with a ‘1’ and excluded from analysis. To account for inter-specific variation in call volume and frequency, consistency for applying ‘background’ designation was approached on a species-specific basis. In cases where vocalizations from the same species were separated by fewer than 5 s, they were included as part of the same annotation, while annotations were split if the gap between vocalizations exceeded 5 s. An assortment of unlabeled sound clips featuring identified and unidentified species were sent to regional experts for secondary verification. Annotations were reviewed repeatedly until unidentified vocalizations made up less than 10% of the total avian vocal activity on a given recording. We made considerable effort to accurately label as many foreground and background vocalizations as possible to improve the value of this dataset for training soundscape-based automated identification algorithms.

**Figure 2:**
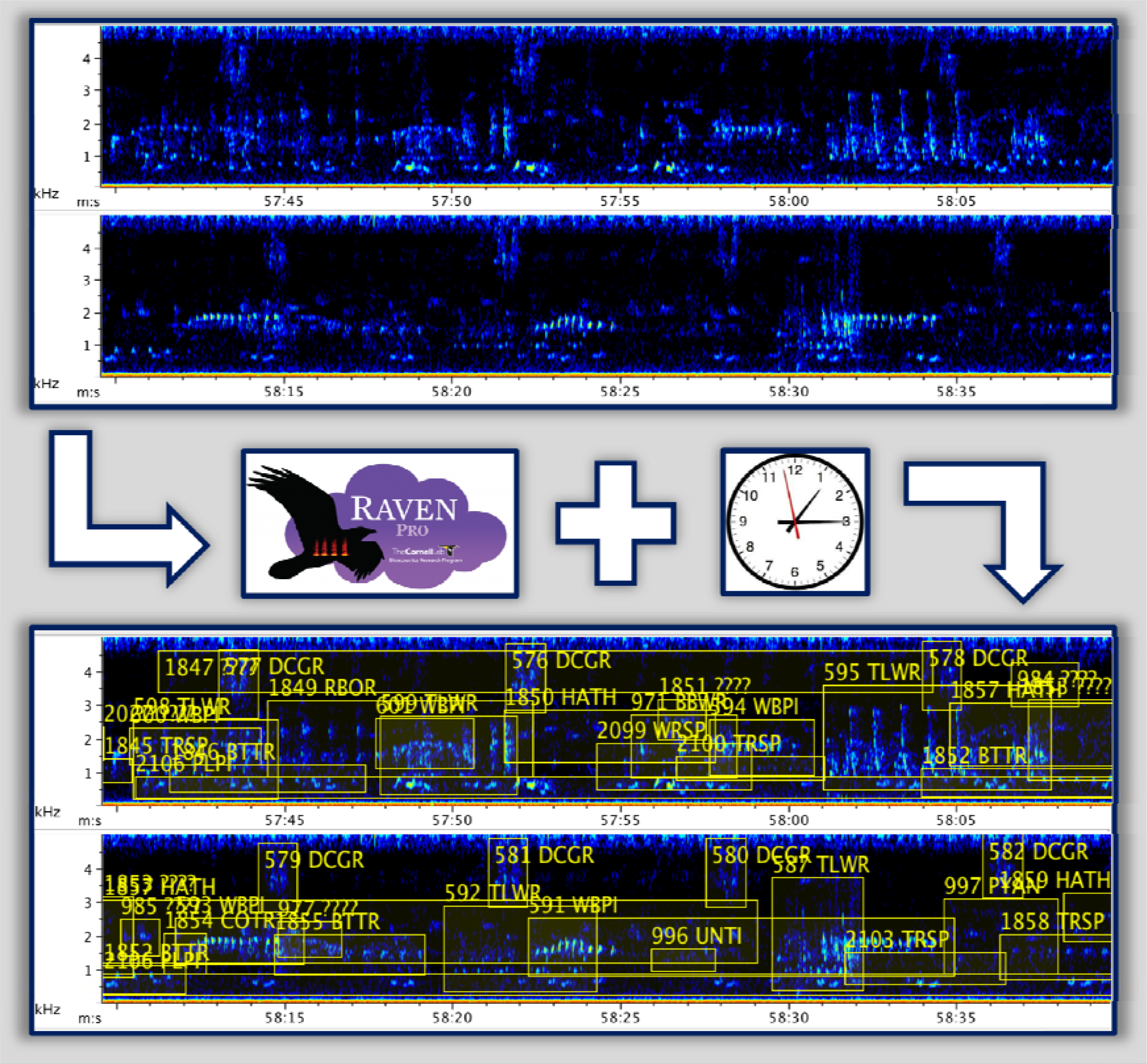
The annotation process. Spectrograms showing one minute of sound from Site B, Day 2, before and after manual annotation in Raven Pro 1.6.

### 2.4 Statistical analyses

To assess fine-scale temporal variation in dawn chorus vocal activity within recordings and across sampling sites and days, we subsampled each 60 min recording at 1 min intervals, totaling 1,080 samples across all sites and days. For each 1 min interval, we calculated species richness (SR), vocal prevalence (VP), and total vocal prevalence (TVP). Species richness refers to the number of unique species detected, while vocal prevalence refers to the number of 10 s intervals featuring a vocalization of a given species. Therefore, a single species could attain a maximum VP of 6 per 1 min interval or 360 per hour recording. Total vocal prevalence (TVP)—a proxy for the total avian vocal activity in a soundscape—refers to the sum of VP for all detected taxa in a given subsample. We assessed the relationship between TVP and SR using a Pearson’s *R* correlation analysis.

Certain taxa were excluded from species richness calculations for data quality reasons: parrots (Psittacidae sp.), because they occurred almost exclusively as calling flyovers, as well as Columbiformes (pigeons and doves), and three congeneric tinamou species, Cinereous Tinamou (*Crypturellus cinereus*), Little Tinamou (*C. soui*), and Bartlett’s Tinamou (*C. bartletti*), which had overlapping vocalizations and were inconsistently identified with confidence. However, these species were included in calculations of TVP, as were vocalizations that were identified to a broader group but not at the species level (e.g., Trogon sp. [Trogonidae sp.]) (For a full list of species and exclusions for SR and TVP calculations, see Supporting Information Appendix S1).

#### 2.4.1 Non-metric multidimensional scaling ordination (NMDS) and ANOVA

We used non-metric multidimensional scaling ordination (NMDS) and two-way repeated-measures ANOVA to assess evidence for significant effects of site-specific differences in community composition on SR and TVP. Substituting VP in place of abundance, we calculated NMDS using the “bray” method, referring to Bray-Curtis dissimilarity, and metaMDS function from the “vegan” package in R (Okansen *et al*. 2019). One advantage of using Bray-Curtis dissimilarity is its ability to account for the relative abundances of different species or taxa (Ricotta & Podani 2017). We plotted NMDS ordinations on two axes and ensured that the output met minimum stress requirements (Kruskal 1964). To compare SR and TVP between sites, days, and recordings, two-way repeated-measures ANOVA followed by Tukey’s HSD tests for multiple comparisons were conducted in R, with site and day as interacting independent variables and the 1-minute time interval as a blocking variable.

#### 2.4.2 Generalized Additive Models (GAMs)

We then constructed a set of Generalized Additive Models (GAMs) fit with Poisson distributions and restricted maximum likelihood estimations to account for the non-linearity of within-recording response curves for SR, TVP, and species-specific VP within the dawn hour. For both the SR and TVP models, we included a smoothing term for the interaction between time of day and recording to produce individual response curves for each recording, and a nested site-day random effects term to account for pseudoreplication. Similarly, to produce species-specific VP response curves across days and sites, we replicated this random effects structure, but removed the interaction for the time interval smoothing term. As the amount of species present at time step *t* directly influences the number of species that are likely to be detected at time step *t* + 1, we accounted for this inherent temporal autocorrelation by fitting both models with a first-order autoregressive covariance structure (AR1; Yang et al. 2012). This decision successfully removed evidence of autocorrelation within model residuals. Because our model structure was biologically informed, we chose to forgo model selection and base inferences and predictions from this structure. We completed all statistical analyses in R (v. 4.2.0; R Core Team 2022) using functions from the “tidyverse” (Wickham et al. 2019) and employing the “mgcv” package to construct GAMs (Wood et al. 2016; Wood 2004, 2011, 2017).

## 3 Results

We identified 127 species over the 18 analyzed dawn hours, seven of which were recorded only as background vocalizations; 17,036 individual bounding-box annotations were drawn in Raven Pro 1.6, of which 94.8% were identified to the species or taxon level. Site A had the lowest average species richness per 1 min interval (4.0; 43 total), followed by Site F (4.4; 42 total), Site D (4.8; 49 total), Site C (4.9; 46 total), Site E (5.3; 55 total), and Site B (6.1; 50 total), which was also the most active site by TVP. Aggregating detections across sampling sites, Day 1 featured 57 species, Day 2 featured 78 species, and Day 3 featured 56 species, with average SR values per 1 min interval of 4.79, 6.18, and 3.81, respectively. At individual sites, the percentage of total vocal prevalence (TVP) that was not identified to a species or taxa group ranged from 4.1% (Site E) to 7.1% (Site B) between sites, and from 5.1% (Day 3) to 5.4% (Day 1) between days.

Of the 120 foreground species detected, only two, Black-faced Antthrush (*Formicarius analis*) and Buff-throated Woodcreeper (*Xiphorhynchus guttatus*), were detected on all 18 recordings, and six more, Hauxwell’s Thrush (*Turdus hauxwelli*), Little Tinamou (*Crypturellus soui*), Thrush-like Wren (*Camplyothynchs turdinus*), Amazonian Motmot (*Momotus momota*), Plumbeous Pigeon (*Patagioenas plumbea*), and Screaming Piha (*Lipaugus cociferans*) were detected on at least 15 recordings. Most species were rare; more than half (*n* = 66, 55%) were detected on 4 or fewer 1-hr recordings, and roughly a quarter (*n* = 27, 23%) on only a single recording (Supporting Information Appendix S2). Regarding total vocal prevalence, the most abundant species were Hauxwell’s Thrush, Black-faced Antthrush, and Thrush-like Wren. TVP was highest at all six sites on 20 January (Day 2) and lowest on 31 January (Day 3) at every site except Site B.

### 3.1 Non-metric multidimensional scaling ordination (NMDS) and ANOVA

Individual sites consistently clustered separately from each other in terms of their species composition, as demonstrated by non-metric multidimensional scaling (NMDS) ordination (Fig. 3). Tukey’s HSD tests indicated that a higher percentage of pairwise comparisons between sites were significantly different (*p* < 0.01) for SR (9/15, 60%) than for TVP (6/15, 40%), and all comparisons between days were significant for both TVP and SR. The rate that recordings from different days exhibited significant pairwise differences (*p* < 0.01) remained essentially identical whether they came from the same site or from different sites: 44.4% (8/18, same site) versus 42.2% (38/90, different sites) for SR and 55.6% (10/18) versus 56.7% (51/90) for TVP. In general, pairwise comparisons of recordings from different days were more likely to differ significantly (*p* < 0.01) than recordings from different sites. For SR, 42.6% of recordings from different days (46/108) differed significantly, compared to 35.6% from different sites (48/135). For TVP, 56.5% of recording pairings from different days (61/108) differed significantly, compared to 45.2% from different sites (61/135). TVP and SR were least likely to differ between recordings that came from different sites on the same day; just 22.2% of these pairwise comparisons (10/45) differed significantly (*p <* 0.01) for each metric. For all Tukey’s HSD pairwise comparisons between sites, days, and individual recordings, see Supporting Information Appendix S3.

**Figure 3:**
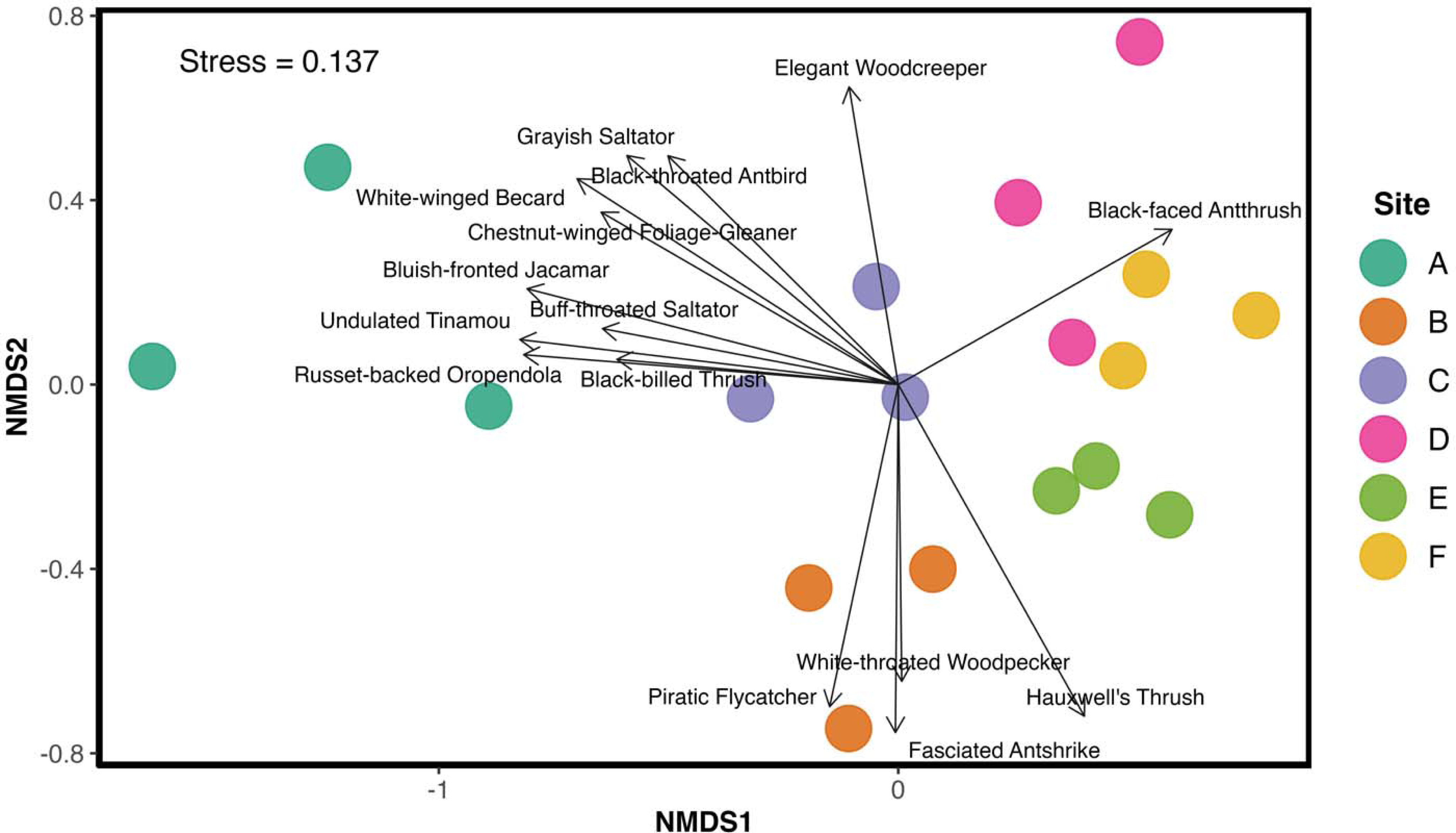
**Non-metric multidimensional scaling (NMDS) ordination** by site using Bray-Curtis dissimilarity, with vocal prevalence in place of abundance. Significantly representative species along the ordination axes (*p <* 0.02) are shown with vectors. For site information, see Fig. 1.

### 3.2 Generalized Additive Models (GAMs)

SR and TVP models accounted for a substantial amount of variation in dawn chorus vocal activity (*R^2^* = 0.64 and *R^2^* = 0.83, respectively, Fig. 4), in which smoothing terms for the interaction between time interval and recording, as well as site-day random intercepts, were often significant (*p* < 0.05; Supporting Information Appendix S4). The effect of time interval and day was generally stronger than the effect of site in explaining variation in observed SR and TVP, and TVP was strongly correlated with SR (*r* = 0.85, *p* < 0.001). We observed that diel variation in avian vocal activity varied by species and taxonomic group, with the VP of different species peaking at different time intervals within the dawn hour (Fig. 5).

**Figure 4:**
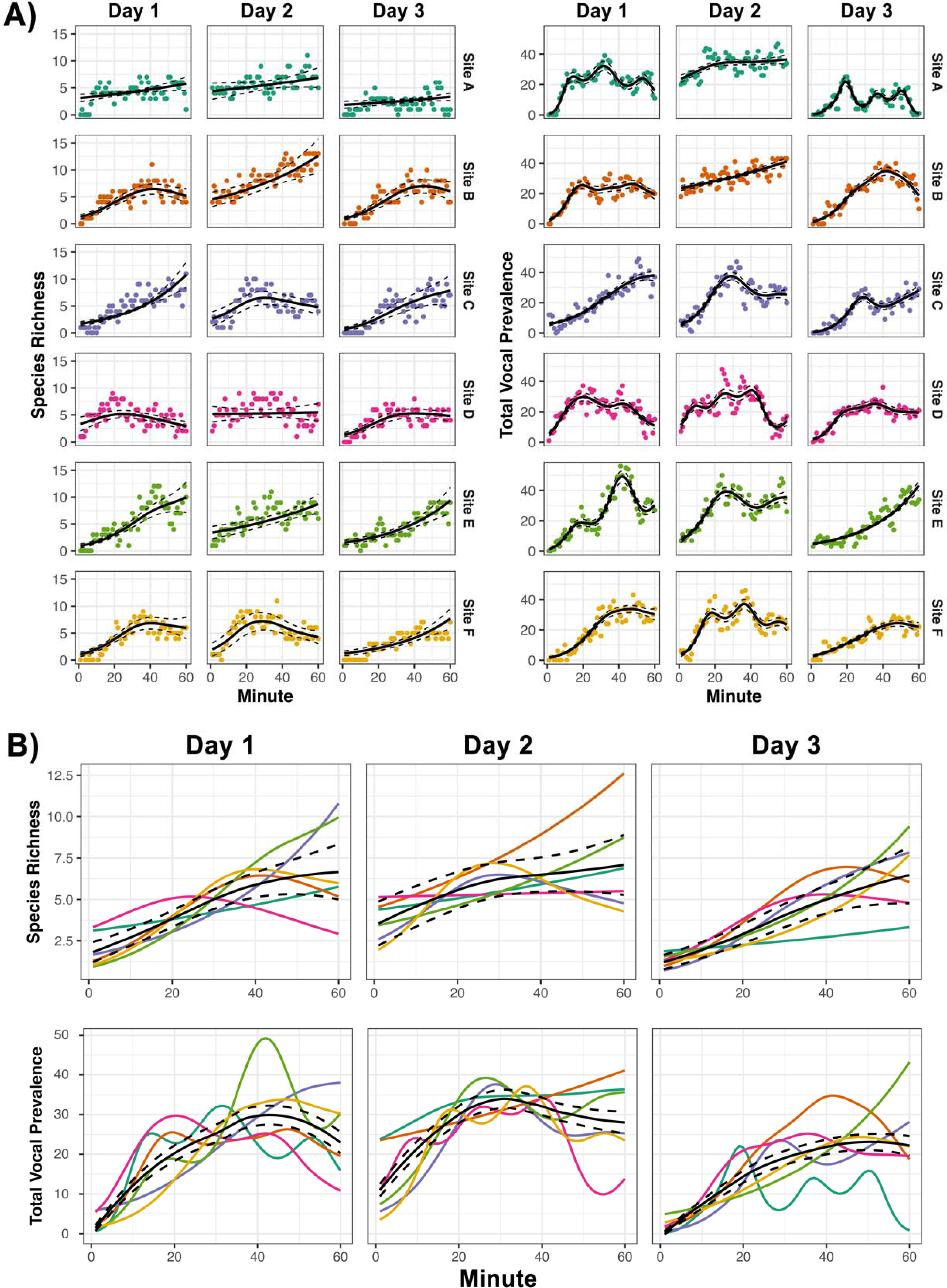
Comparisons of (A) species richness and total vocal prevalence detected by time of day across days and sites overlaid with Generalized Additive Model (GAM) prediction curves ± 95% confidence intervals, along with (B) site-specific GAM prediction curves overlaid with day-specific mean GAM prediction curves ± 95% confidence intervals of species richness and vocal prevalence across days.

**Figure 5:**
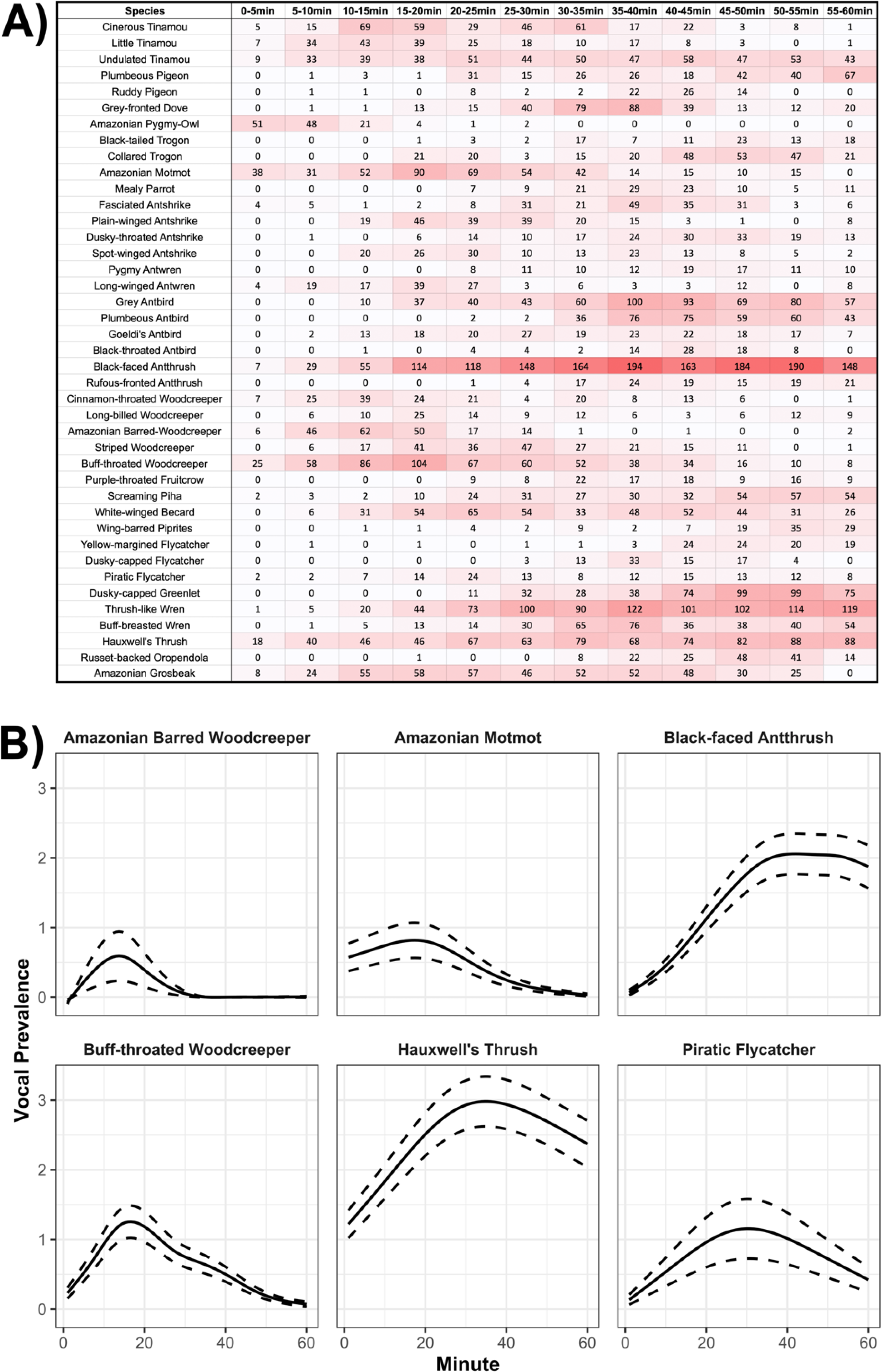
Species-level diel variation in vocal prevalence within the dawn hour. (A) Diel variation in vocal prevalence by 5 min interval for species with total vocal prevalence > 75 from all sites and days, and (B) model-generated prediction curves ± 95% confidence intervals of species-specific vocal prevalence within the dawn hour for six species (B).

## 4 Discussion

We used simultaneous passive acoustic monitoring to quantify temporal variation in the vocal activity level and observed species richness of an Amazonian bird community. In line with our primary hypothesis, the magnitude of this temporal variation is large enough to potentially mask, or even supersede, observed differences in species richness and vocal activity levels between sites and habitat types—complicating the interpretation of point count studies and other non-simultaneous survey methods that fail to control for temporal variables. While temporal variation in avian activity over the course of the morning is well-documented in the tropics, the limitations of traditional non-simultaneous survey methods mean that point count studies on temporal effects in these systems offer only broad conclusions, typically relating to maximum temporal resolutions of one hour or more (Blake 1992, Lynch 1995, Woltmann 2005, Esquivel & Peris 2008). Fine-scale diel variation in the vocal activity and detectability of Amazonian birds has been referenced anecdotally for decades (Parker 1991). However, thanks to the development of autonomous recording units, which allow for considerably easier repeat sampling than traditional methods, we are beginning to understand more about these effects at higher temporal resolutions (Gil & Llusia 2020, Blake 2021, Metcalf *et al*. 2021, Pérez-Granados & Schuchmann 2023). While prior temporal effects studies have focused primarily on diel and seasonal temporal variation, to our knowledge, this is the first study to provide evidence that day-to-day temporal variation in the vocal activity and detectability of Amazonian bird communities may be strong enough to obscure statistically significant differences between sites, in the absence of simultaneous data collection.

The variable diel patterns in avian vocal activity that we observed between species and taxonomic groups (Fig. 5) mean that visiting two sites with the same overall SR or TVP at different times in the morning could result in considerably different observations of community composition, even when both visits occur within the same hour. This species-to-species diel variation in vocal activity and detectability may partially or entirely explain why the diel variation patterns in SR and TVP that we documented within the dawn hour were spatially heterogeneous. Consistent with the findings of Rodriguez *et al*. (2014), these patterns varied per-site, even between sites in the same general habitat type and superficial condition. Because sites with differing communities should be expected to exhibit differing patterns of diel variation in overall vocal activity, the differing patterns of diel variation in TVP and SR that we documented between sites are plausibly the result of the spatial heterogeneity of avian community composition (Fig. 3).

The spatial heterogeneity that we observed in contiguous lowland forest with superficially similar conditions may have been the result of cryptic habitat diversity. Though sometimes misrepresented as homogenous, Amazonian habitats are highly heterogeneous (Pires & Prance 1985, Tuomisto *et al*. 1995), and important elements of their structural diversity remain poorly understood (Macía 2011, Borges 2013). Interactions between habitat features and avian community structure in the Amazon are extremely complex and are influenced by variables including soil type (Borges 2013), riverine sediment concentration (Laranjeiras *et al*. 2021), the condition of surrounding habitat (Latta *et al*. 2011, Laurence *et al*. 2011, Woltmann 2012, Laurence *et al*. 2012, Wolfe *et al*. 2015, Barlow 2016, Menger *et al*. 2017, Hernández-Palma & Stouffer 2018), the presence of large animals (Redford 1992, Estes *et al*. 2011), and climate change (Blake & Loiselle 2015, Stouffer *et al*. 2020). As a result, Amazonian habitat assessments, which generally rely on features discernable by field observers, can be overly simplistic (Milliken *et al*. 2010), and we avoided incorporating them in our analysis for this reason. The complexity of Amazonian systems increases the importance of controlling for temporal variables, which may obscure these effects.

While this study was not designed to serve as a comprehensive census of the avian community at ITRA, our observations were generally consistent with those documented by previous census efforts in Madre de Dios’s *várzea* forests (Terborgh *et al*. 1990, Martínez *et al*. 2023), namely that most species were rare, with a high proportion of TVP contributed by only a few species, and that sites were compositionally heterogeneous. The observation that the site in edge habitat (Site A) had the lowest average species richness per 1-min interval (4.03; 43 total), followed by Site F (4.41; 42 total), in degraded forest outside of the reserve boundaries, is consistent with the results of research focused on edge effects in Amazonia, which are known to pervasively impact Amazonian bird communities (Terborgh *et al*. 1990, Laurence 2004, Barlow *et al*. 2006, Broadbent *et al*. 2008, Laurence *et al*. 2011, Moura *et al*. 2016, Luther *et al*. 2020, Stouffer 2020). This habitat usage pattern differs from observed dynamics in temperate systems, where avian diversity tends to be higher in edge habitats than forest interiors (Baldi 1996, Lindell *et al*. 2007). Most annotations occurred in the frequency range of 0.5–5 kHz, suggesting possible signal masking from insects, which primarily occur between 4–12 kHz (Hart *et al*. 2015, Metcalf *et al*. 2020).

The temporal variation in avian vocal activity levels and observed species richness that we documented between days was greater and more consistent than we expected, generally influencing SR and TVP more than differences between sites. Supporting the idea that temporal effects may overwhelm statistically significant differences between sites, sampling the same site on different days was more than twice as likely to result in significantly differing measures of SR and TVP than sampling different sites on the same day, even though most site-to-site comparisons indicated significant differences in SR or TVP. That temporal variation was strongest for TVP is relevant for point counts because aural detection probability for a field observer is a direct function of VP, which reflects the probability that a species vocalizes during a given 10 s window. Standard duration point counts should have higher rates of false absences on days with lower TVP, even with static species availability, because of the reduction in time windows where species vocalize.

We found that TVP was a useful metric for quantifying avian vocal activity, and believe that it is worthy of further study and use. TVP is a more stable indicator of overall activity levels than raw call count, total number of annotations, or total annotation length because it is robust to natural differences in vocalization patterns across species. When annotation boxes are split, if the gap between vocalizations exceeds a set time interval (5 s in this study), alternative activity metrics are dependent on the innate vocalization rate of a given species. For example, Hauxwell’s Thrush (HATH) and Black-faced Antthrush (BFAT) were two of the most common species in this dataset but feature different vocalization styles. While HATHs often take only short breaks of < 5 s between song bouts in the morning, BFATs sing with gaps between song bouts that are generally greater than 5 s. A HATH singing throughout the morning may only result in a small number of different annotation boxes, but a large total annotation length that incorporates the time in-between song bouts, while a BFAT singing throughout the morning can feature a very high number of total annotations, but will not include the gaps between song phrases because they exceed 5 s in length. In our data, BFAT had a total annotation length of 4,456 seconds across 1,500 individual bounding boxes, while HATH had a total annotation length of 25,308 seconds that constituted just 675 total bounding boxes. Using TVP eliminates this type of artificial species-to-species variability in vocal activity estimates. As a result, TVP can improve comparisons of annotated bioacoustic datasets that employ slightly different annotation protocols and can reduce manual analysis time by eliminating the need to measure the distance between vocalizations. While metrics that require species-level ID are highly contingent on observer skill level, driving inconsistencies in Neotropical bird surveys (Robinson *et al*. 2018), TVP supports using broader taxonomic groups for challenging identifications, potentially lowering time and experience barriers for researchers. We found that TVP correlated strongly with SR (*r* = 0.85, *p* < 0.001), meaning that it may be a viable proxy for species richness, even without requiring species IDs, and should be easier to generate with automated approaches than metrics that require species-level IDs. Because TVP essentially represents soundscape abundance, it can also enable the calculation of abundance-based indices like Bray-Curtis dissimilarity and Shannon diversity with acoustic data.

Based on our collective results, we suggest that bioacoustic data should be collected to complement traditional avian surveys whenever possible. In addition to the short-term benefits of quantifying diel and day-to-day temporal variation, even small bioacoustic datasets serve as ecological time capsules (Sugai & Llusia 2019), which are particularly important in an era of rapid global change. ARUs can generate greater data volume than traditional methods without meaningfully increasing field time or cost (Hobson *et al*. 2002, Acevedo & Villanueva-Rivera 2006, Tegeler *et al*. 2012). While manually analyzing acoustic data can be time-intensive, limiting its utility, acoustic indices (Jorge *et al*. 2018, Metcalf *et al*. 2020) and automated identification programs like BirdNET (Kahl *et al*. 2021) are continuously improving and could have a transformative effect on global ornithological research (Pérez-Granados 2023). Due to their relative rarity, fully-annotated tropical soundscapes are critical for developing these automated programs, and this dataset has already been used for this purpose (Kahl *et al*. 2020). Studies focused exclusively on birds can come at the expense of developing knowledge of other groups (Gardner *et al*. 2008), but bioacoustic data collected for ornithological research often contains vocalizations of non-target taxa and other soundscape elements, enabling more holistic biodiversity research (Newson *et al*. 2017). To help advance these efforts, we encourage researchers collecting bioacoustic data to make use of open-access repositories like Zenodo (European Organization For Nuclear Research & OpenAIRE 2013) to host their datasets, as we have (Hopping *et al*. 2022).

To conclude, we found that simultaneous passive acoustic monitoring coupled with manual annotation revealed significant diel and day-to-day variation in the vocal activity and observed species richness of an Amazonian bird community, an ecological pattern that would be difficult to ascertain using traditional field methods. The magnitude of this temporal variation was large enough that it could mask meaningful differences between sites if temporal bias is not sufficiently accounted for in study designs. This research provides a case study for using passive acoustic monitoring to quantify temporal variation in tropical ornithological surveys, even with a relatively small sample of sites and days. Future studies could help explain the mechanisms for this variation and improve methods for processing and interpreting large volumes of acoustic data in complex systems.

## Authors’ Contributions

W.A.H. and H.K. designed the methodology and secured the funding to conduct the study; W.A.H. and C.J.S. analyzed the data; W.A.H. and N.H.C. collected and annotated the recordings, and W.A.H. led the writing of the manuscript. All authors contributed critically to the drafts and gave final approval for publication. The Spanish abstract was translated by N.H.C.

## Acknowledgements

We would like to thank the Inkaterra Association (ITA) staff for providing logistical support and excellent field station facilities, particularly Dennis Osorio and Kevin Jiménez Gonzales, who helped set up recorders. John Fitzpatrick, Fernando Angulo, Will Sweet, Ken Rosenberg, and Alex Wiebe helped identify unknown vocalizations, and Richard Fuller, Ken Rosenberg, Andrew Farnsworth, Connor Wood, and several anonymous reviewers provided helpful comments on the drafts.

## Intellectual Contributions

Our study involved collaboration with multiple Peruvian NGOs, particularly the Inkaterra Association (ITA), in both Lima and Madre de Dios. While in the field, we taught local researchers how to set up and maintain Swift recorders, how to annotate recordings in Raven Pro, and then left 9 recorders with them for their own use. We shared a copy of all collected data and our results with ITA, and one of ‘ITA’s local researchers (Noe Huaraca-Charca) is a coauthor on this paper.

## Data Availability Statement

All software curated for this research are archived and available at https://github.com/csayers2/Inkaterra-ARU. The raw acoustic data and annotations referenced in this study can be found at https://zenodo.org/record/7079124 (DOI: 10.5281/zenodo.7079124).

