## Supplementary figures and images for "Simultaneous passive acoustic monitoring uncovers evidence of potentially overlooked temporal variation in an Amazonian bird community"

### Supporting Information Appendix S2

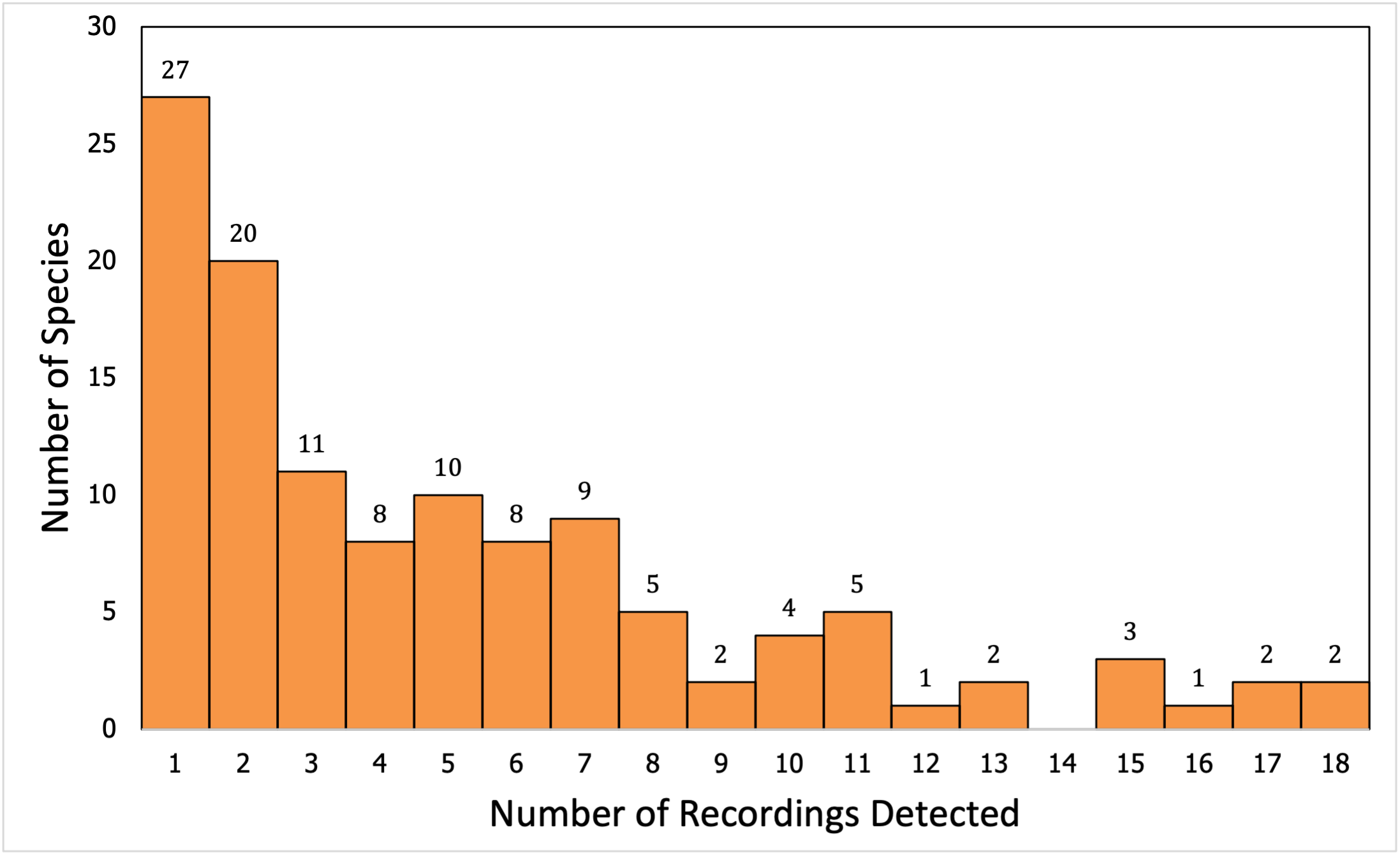
