## Supporting Information Appendix S3 for "Simultaneous passive acoustic monitoring uncovers evidence of potentially overlooked temporal variation in an Amazonian bird community"

**Supporting Information Appendix S3: Tukey’s HSD comparisons between sites, days, and recordings.**

**Total Vocal Prevalence:**

```
  Tukey multiple comparisons of means
    95% family-wise confidence level

Fit: aov(formula = TVP ~ Site * Day + Time.Window, data = timesplits)

$Site
                     diff         lwr        upr     p adj
Site B-Site A  3.25555556  0.72392237  5.7871887 0.0034463
Site C-Site A -0.50000000 -3.03163318  2.0316332 0.9932802
Site D-Site A -0.06666667 -2.59829985  2.4649665 0.9999997
Site E-Site A  2.60555556  0.07392237  5.1371887 0.0394201
Site F-Site A -0.82222222 -3.35385540  1.7094110 0.9395245
Site C-Site B -3.75555556 -6.28718874 -1.2239224 0.0003565
Site D-Site B -3.32222222 -5.85385540 -0.7905890 0.0025956
Site E-Site B -0.65000000 -3.18163318  1.8816332 0.9778433
Site F-Site B -4.07777778 -6.60941096 -1.5461446 0.0000698
Site D-Site C  0.43333333 -2.09829985  2.9649665 0.9965690
Site E-Site C  3.10555556  0.57392237  5.6371887 0.0063785
Site F-Site C -0.32222222 -2.85385540  2.2094110 0.9991728
Site E-Site D  2.67222222  0.14058904  5.2038554 0.0315856
Site F-Site D -0.75555556 -3.28718874  1.7760776 0.9574817
Site F-Site E -3.42777778 -5.95941096 -0.8961446 0.0016370

$Day
                  diff        lwr        upr p adj
Day 2-Day 1   6.108333   4.636687   7.579980     0
Day 3-Day 1  -5.430556  -6.902202  -3.958909     0
Day 3-Day 2 -11.538889 -13.010535 -10.067243     0

$`Site:Day`
                                  diff          lwr           upr     p adj
Site B:Day 1-Site A:Day 1  -0.58333333  -5.95505596   4.788389294 1.0000000
Site C:Day 1-Site A:Day 1   0.93333333  -4.43838929   6.305055961 1.0000000
Site D:Day 1-Site A:Day 1   0.66666667  -4.70505596   6.038389294 1.0000000
Site E:Day 1-Site A:Day 1   3.71666667  -1.65505596   9.088389294 0.5928211
Site F:Day 1-Site A:Day 1  -0.48333333  -5.85505596   4.888389294 1.0000000
Site A:Day 2-Site A:Day 1  12.11666667   6.74494404  17.488389294 0.0000000
Site B:Day 2-Site A:Day 1  10.63333333   5.26161071  16.005055961 0.0000000
Site C:Day 2-Site A:Day 1   3.65000000  -1.72172263   9.021722627 0.6261087
Site D:Day 2-Site A:Day 1   2.55000000  -2.82172263   7.921722627 0.9733103
Site E:Day 2-Site A:Day 1   8.00000000   2.62827737  13.371722627 0.0000338
Site F:Day 2-Site A:Day 1   3.95000000  -1.42172263   9.321722627 0.4757448
Site A:Day 3-Site A:Day 1 -11.55000000 -16.92172263  -6.178277373 0.0000000
Site B:Day 3-Site A:Day 1   0.28333333  -5.08838929   5.655055961 1.0000000
Site C:Day 3-Site A:Day 1  -5.51666667 -10.88838929  -0.144944039 0.0367471
Site D:Day 3-Site A:Day 1  -2.85000000  -8.22172263   2.521722627 0.9271388
Site E:Day 3-Site A:Day 1  -3.33333333  -8.70505596   2.038389294 0.7729851
Site F:Day 3-Site A:Day 1  -5.36666667 -10.73838929   0.005055961 0.0505282
Site C:Day 1-Site B:Day 1   1.51666667  -3.85505596   6.888389294 0.9999522
Site D:Day 1-Site B:Day 1   1.25000000  -4.12172263   6.621722627 0.9999971
Site E:Day 1-Site B:Day 1   4.30000000  -1.07172263   9.671722627 0.3146097
Site F:Day 1-Site B:Day 1   0.10000000  -5.27172263   5.471722627 1.0000000
Site A:Day 2-Site B:Day 1  12.70000000   7.32827737  18.071722627 0.0000000
Site B:Day 2-Site B:Day 1  11.21666667   5.84494404  16.588389294 0.0000000
Site C:Day 2-Site B:Day 1   4.23333333  -1.13838929   9.605055961 0.3429943
Site D:Day 2-Site B:Day 1   3.13333333  -2.23838929   8.505055961 0.8490498
Site E:Day 2-Site B:Day 1   8.58333333   3.21161071  13.955055961 0.0000044
Site F:Day 2-Site B:Day 1   4.53333333  -0.83838929   9.905055961 0.2262466
Site A:Day 3-Site B:Day 1 -10.96666667 -16.33838929  -5.594944039 0.0000000
Site B:Day 3-Site B:Day 1   0.86666667  -4.50505596   6.238389294 1.0000000
Site C:Day 3-Site B:Day 1  -4.93333333 -10.30505596   0.438389294 0.1169598
Site D:Day 3-Site B:Day 1  -2.26666667  -7.63838929   3.105055961 0.9921330
Site E:Day 3-Site B:Day 1  -2.75000000  -8.12172263   2.621722627 0.9463233
Site F:Day 3-Site B:Day 1  -4.78333333 -10.15505596   0.588389294 0.1518315
Site D:Day 1-Site C:Day 1  -0.26666667  -5.63838929   5.105055961 1.0000000
Site E:Day 1-Site C:Day 1   2.78333333  -2.58838929   8.155055961 0.9403857
Site F:Day 1-Site C:Day 1  -1.41666667  -6.78838929   3.955055961 0.9999819
Site A:Day 2-Site C:Day 1  11.18333333   5.81161071  16.555055961 0.0000000
Site B:Day 2-Site C:Day 1   9.70000000   4.32827737  15.071722627 0.0000001
Site C:Day 2-Site C:Day 1   2.71666667  -2.65505596   8.088389294 0.9518220
Site D:Day 2-Site C:Day 1   1.61666667  -3.75505596   6.988389294 0.9998839
Site E:Day 2-Site C:Day 1   7.06666667   1.69494404  12.438389294 0.0006612
Site F:Day 2-Site C:Day 1   3.01666667  -2.35505596   8.388389294 0.8855601
Site A:Day 3-Site C:Day 1 -12.48333333 -17.85505596  -7.111610706 0.0000000
Site B:Day 3-Site C:Day 1  -0.65000000  -6.02172263   4.721722627 1.0000000
Site C:Day 3-Site C:Day 1  -6.45000000 -11.82172263  -1.078277373 0.0037923
Site D:Day 3-Site C:Day 1  -3.78333333  -9.15505596   1.588389294 0.5592586
Site E:Day 3-Site C:Day 1  -4.26666667  -9.63838929   1.105055961 0.3286405
Site F:Day 3-Site C:Day 1  -6.30000000 -11.67172263  -0.928277373 0.0056383
Site E:Day 1-Site D:Day 1   3.05000000  -2.32172263   8.421722627 0.8757502
Site F:Day 1-Site D:Day 1  -1.15000000  -6.52172263   4.221722627 0.9999992
Site A:Day 2-Site D:Day 1  11.45000000   6.07827737  16.821722627 0.0000000
Site B:Day 2-Site D:Day 1   9.96666667   4.59494404  15.338389294 0.0000000
Site C:Day 2-Site D:Day 1   2.98333333  -2.38838929   8.355055961 0.8948689
Site D:Day 2-Site D:Day 1   1.88333333  -3.48838929   7.255055961 0.9991274
Site E:Day 2-Site D:Day 1   7.33333333   1.96161071  12.705055961 0.0002940
Site F:Day 2-Site D:Day 1   3.28333333  -2.08838929   8.655055961 0.7934837
Site A:Day 3-Site D:Day 1 -12.21666667 -17.58838929  -6.844944039 0.0000000
Site B:Day 3-Site D:Day 1  -0.38333333  -5.75505596   4.988389294 1.0000000
Site C:Day 3-Site D:Day 1  -6.18333333 -11.55505596  -0.811610706 0.0076142
Site D:Day 3-Site D:Day 1  -3.51666667  -8.88838929   1.855055961 0.6908651
Site E:Day 3-Site D:Day 1  -4.00000000  -9.37172263   1.371722627 0.4511957
Site F:Day 3-Site D:Day 1  -6.03333333 -11.40505596  -0.661610706 0.0110865
Site F:Day 1-Site E:Day 1  -4.20000000  -9.57172263   1.171722627 0.3576577
Site A:Day 2-Site E:Day 1   8.40000000   3.02827737  13.771722627 0.0000085
Site B:Day 2-Site E:Day 1   6.91666667   1.54494404  12.288389294 0.0010283
Site C:Day 2-Site E:Day 1  -0.06666667  -5.43838929   5.305055961 1.0000000
Site D:Day 2-Site E:Day 1  -1.16666667  -6.53838929   4.205055961 0.9999990
Site E:Day 2-Site E:Day 1   4.28333333  -1.08838929   9.655055961 0.3215839
Site F:Day 2-Site E:Day 1   0.23333333  -5.13838929   5.605055961 1.0000000
Site A:Day 3-Site E:Day 1 -15.26666667 -20.63838929  -9.894944039 0.0000000
Site B:Day 3-Site E:Day 1  -3.43333333  -8.80505596   1.938389294 0.7294115
Site C:Day 3-Site E:Day 1  -9.23333333 -14.60505596  -3.861610706 0.0000004
Site D:Day 3-Site E:Day 1  -6.56666667 -11.93838929  -1.194944039 0.0027640
Site E:Day 3-Site E:Day 1  -7.05000000 -12.42172263  -1.678277373 0.0006948
Site F:Day 3-Site E:Day 1  -9.08333333 -14.45505596  -3.711610706 0.0000007
Site A:Day 2-Site F:Day 1  12.60000000   7.22827737  17.971722627 0.0000000
Site B:Day 2-Site F:Day 1  11.11666667   5.74494404  16.488389294 0.0000000
Site C:Day 2-Site F:Day 1   4.13333333  -1.23838929   9.505055961 0.3878517
Site D:Day 2-Site F:Day 1   3.03333333  -2.33838929   8.405055961 0.8807178
Site E:Day 2-Site F:Day 1   8.48333333   3.11161071  13.855055961 0.0000063
Site F:Day 2-Site F:Day 1   4.43333333  -0.93838929   9.805055961 0.2619307
Site A:Day 3-Site F:Day 1 -11.06666667 -16.43838929  -5.694944039 0.0000000
Site B:Day 3-Site F:Day 1   0.76666667  -4.60505596   6.138389294 1.0000000
Site C:Day 3-Site F:Day 1  -5.03333333 -10.40505596   0.338389294 0.0974360
Site D:Day 3-Site F:Day 1  -2.36666667  -7.73838929   3.005055961 0.9874716
Site E:Day 3-Site F:Day 1  -2.85000000  -8.22172263   2.521722627 0.9271388
Site F:Day 3-Site F:Day 1  -4.88333333 -10.25505596   0.488389294 0.1278126
Site B:Day 2-Site A:Day 2  -1.48333333  -6.85505596   3.888389294 0.9999651
Site C:Day 2-Site A:Day 2  -8.46666667 -13.83838929  -3.094944039 0.0000067
Site D:Day 2-Site A:Day 2  -9.56666667 -14.93838929  -4.194944039 0.0000001
Site E:Day 2-Site A:Day 2  -4.11666667  -9.48838929   1.255055961 0.3955685
Site F:Day 2-Site A:Day 2  -8.16666667 -13.53838929  -2.794944039 0.0000191
Site A:Day 3-Site A:Day 2 -23.66666667 -29.03838929 -18.294944039 0.0000000
Site B:Day 3-Site A:Day 2 -11.83333333 -17.20505596  -6.461610706 0.0000000
Site C:Day 3-Site A:Day 2 -17.63333333 -23.00505596 -12.261610706 0.0000000
Site D:Day 3-Site A:Day 2 -14.96666667 -20.33838929  -9.594944039 0.0000000
Site E:Day 3-Site A:Day 2 -15.45000000 -20.82172263 -10.078277373 0.0000000
Site F:Day 3-Site A:Day 2 -17.48333333 -22.85505596 -12.111610706 0.0000000
Site C:Day 2-Site B:Day 2  -6.98333333 -12.35505596  -1.611610706 0.0008461
Site D:Day 2-Site B:Day 2  -8.08333333 -13.45505596  -2.711610706 0.0000255
Site E:Day 2-Site B:Day 2  -2.63333333  -8.00505596   2.738389294 0.9637528
Site F:Day 2-Site B:Day 2  -6.68333333 -12.05505596  -1.311610706 0.0020010
Site A:Day 3-Site B:Day 2 -22.18333333 -27.55505596 -16.811610706 0.0000000
Site B:Day 3-Site B:Day 2 -10.35000000 -15.72172263  -4.978277373 0.0000000
Site C:Day 3-Site B:Day 2 -16.15000000 -21.52172263 -10.778277373 0.0000000
Site D:Day 3-Site B:Day 2 -13.48333333 -18.85505596  -8.111610706 0.0000000
Site E:Day 3-Site B:Day 2 -13.96666667 -19.33838929  -8.594944039 0.0000000
Site F:Day 3-Site B:Day 2 -16.00000000 -21.37172263 -10.628277373 0.0000000
Site D:Day 2-Site C:Day 2  -1.10000000  -6.47172263   4.271722627 0.9999996
Site E:Day 2-Site C:Day 2   4.35000000  -1.02172263   9.721722627 0.2941945
Site F:Day 2-Site C:Day 2   0.30000000  -5.07172263   5.671722627 1.0000000
Site A:Day 3-Site C:Day 2 -15.20000000 -20.57172263  -9.828277373 0.0000000
Site B:Day 3-Site C:Day 2  -3.36666667  -8.73838929   2.005055961 0.7588236
Site C:Day 3-Site C:Day 2  -9.16666667 -14.53838929  -3.794944039 0.0000005
Site D:Day 3-Site C:Day 2  -6.50000000 -11.87172263  -1.128277373 0.0033143
Site E:Day 3-Site C:Day 2  -6.98333333 -12.35505596  -1.611610706 0.0008461
Site F:Day 3-Site C:Day 2  -9.01666667 -14.38838929  -3.644944039 0.0000009
Site E:Day 2-Site D:Day 2   5.45000000   0.07827737  10.821722627 0.0424067
Site F:Day 2-Site D:Day 2   1.40000000  -3.97172263   6.771722627 0.9999848
Site A:Day 3-Site D:Day 2 -14.10000000 -19.47172263  -8.728277373 0.0000000
Site B:Day 3-Site D:Day 2  -2.26666667  -7.63838929   3.105055961 0.9921330
Site C:Day 3-Site D:Day 2  -8.06666667 -13.43838929  -2.694944039 0.0000270
Site D:Day 3-Site D:Day 2  -5.40000000 -10.77172263  -0.028277373 0.0471323
Site E:Day 3-Site D:Day 2  -5.88333333 -11.25505596  -0.511610706 0.0159464
Site F:Day 3-Site D:Day 2  -7.91666667 -13.28838929  -2.544944039 0.0000448
Site F:Day 2-Site E:Day 2  -4.05000000  -9.42172263   1.321722627 0.4270372
Site A:Day 3-Site E:Day 2 -19.55000000 -24.92172263 -14.178277373 0.0000000
Site B:Day 3-Site E:Day 2  -7.71666667 -13.08838929  -2.344944039 0.0000868
Site C:Day 3-Site E:Day 2 -13.51666667 -18.88838929  -8.144944039 0.0000000
Site D:Day 3-Site E:Day 2 -10.85000000 -16.22172263  -5.478277373 0.0000000
Site E:Day 3-Site E:Day 2 -11.33333333 -16.70505596  -5.961610706 0.0000000
Site F:Day 3-Site E:Day 2 -13.36666667 -18.73838929  -7.994944039 0.0000000
Site A:Day 3-Site F:Day 2 -15.50000000 -20.87172263 -10.128277373 0.0000000
Site B:Day 3-Site F:Day 2  -3.66666667  -9.03838929   1.705055961 0.6178262
Site C:Day 3-Site F:Day 2  -9.46666667 -14.83838929  -4.094944039 0.0000002
Site D:Day 3-Site F:Day 2  -6.80000000 -12.17172263  -1.428277373 0.0014391
Site E:Day 3-Site F:Day 2  -7.28333333 -12.65505596  -1.911610706 0.0003431
Site F:Day 3-Site F:Day 2  -9.31666667 -14.68838929  -3.944944039 0.0000003
Site B:Day 3-Site A:Day 3  11.83333333   6.46161071  17.205055961 0.0000000
Site C:Day 3-Site A:Day 3   6.03333333   0.66161071  11.405055961 0.0110865
Site D:Day 3-Site A:Day 3   8.70000000   3.32827737  14.071722627 0.0000029
Site E:Day 3-Site A:Day 3   8.21666667   2.84494404  13.588389294 0.0000161
Site F:Day 3-Site A:Day 3   6.18333333   0.81161071  11.555055961 0.0076142
Site C:Day 3-Site B:Day 3  -5.80000000 -11.17172263  -0.428277373 0.0194097
Site D:Day 3-Site B:Day 3  -3.13333333  -8.50505596   2.238389294 0.8490498
Site E:Day 3-Site B:Day 3  -3.61666667  -8.98838929   1.755055961 0.6425715
Site F:Day 3-Site B:Day 3  -5.65000000 -11.02172263  -0.278277373 0.0273728
Site D:Day 3-Site C:Day 3   2.66666667  -2.70505596   8.038389294 0.9592812
Site E:Day 3-Site C:Day 3   2.18333333  -3.18838929   7.555055961 0.9948266
Site F:Day 3-Site C:Day 3   0.15000000  -5.22172263   5.521722627 1.0000000
Site E:Day 3-Site D:Day 3  -0.48333333  -5.85505596   4.888389294 1.0000000
Site F:Day 3-Site D:Day 3  -2.51666667  -7.88838929   2.855055961 0.9765352
Site F:Day 3-Site E:Day 3  -2.03333333  -7.40505596   3.338389294 0.9977515
```

**Species Richness:**

```
Tukey multiple comparisons of means
    95% family-wise confidence level

Fit: aov(formula = SR ~ Site * Day + Time.Window, data = timesplits)

$Site
                    diff         lwr        upr     p adj
Site B-Site A  2.0611111  1.46670834  2.6555139 0.0000000
Site C-Site A  0.9000000  0.30559723  1.4944028 0.0002435
Site D-Site A  0.7722222  0.17781945  1.3666250 0.0029918
Site E-Site A  1.2722222  0.67781945  1.8666250 0.0000000
Site F-Site A  0.3833333 -0.21106943  0.9777361 0.4396222
Site C-Site B -1.1611111 -1.75551388 -0.5667083 0.0000005
Site D-Site B -1.2888889 -1.88329166 -0.6944861 0.0000000
Site E-Site B -0.7888889 -1.38329166 -0.1944861 0.0022049
Site F-Site B -1.6777778 -2.27218055 -1.0833750 0.0000000
Site D-Site C -0.1277778 -0.72218055  0.4666250 0.9900574
Site E-Site C  0.3722222 -0.22218055  0.9666250 0.4740810
Site F-Site C -0.5166667 -1.11106943  0.0777361 0.1304475
Site E-Site D  0.5000000 -0.09440277  1.0944028 0.1565357
Site F-Site D -0.3888889 -0.98329166  0.2055139 0.4227030
Site F-Site E -0.8888889 -1.48329166 -0.2944861 0.0003074

$Day
                 diff       lwr        upr p adj
Day 2-Day 1  1.386111  1.040583  1.7316393     0
Day 3-Day 1 -0.975000 -1.320528 -0.6294718     0
Day 3-Day 2 -2.361111 -2.706639 -2.0155829     0

$`Site:Day`
                                   diff         lwr         upr     p adj
Site B:Day 1-Site A:Day 1  8.500000e-01 -0.41122806  2.11122806 0.6408077
Site C:Day 1-Site A:Day 1  9.833333e-01 -0.27789472  2.24456139 0.3629488
Site D:Day 1-Site A:Day 1  5.666667e-01 -0.69456139  1.82789472 0.9846262
Site E:Day 1-Site A:Day 1  1.316667e+00  0.05543861  2.57789472 0.0300785
Site F:Day 1-Site A:Day 1  3.166667e-01 -0.94456139  1.57789472 0.9999912
Site A:Day 2-Site A:Day 1  1.716667e+00  0.45543861  2.97789472 0.0003146
Site B:Day 2-Site A:Day 1  4.333333e+00  3.07210528  5.59456139 0.0000000
Site C:Day 2-Site A:Day 1  1.350000e+00  0.08877194  2.61122806 0.0218071
Site D:Day 2-Site A:Day 1  1.483333e+00  0.22210528  2.74456139 0.0053838
Site E:Day 2-Site A:Day 1  2.033333e+00  0.77210528  3.29456139 0.0000033
Site F:Day 2-Site A:Day 1  1.433333e+00  0.17210528  2.69456139 0.0092848
Site A:Day 3-Site A:Day 1 -1.983333e+00 -3.24456139 -0.72210528 0.0000071
Site B:Day 3-Site A:Day 1  7.333333e-01 -0.52789472  1.99456139 0.8524089
Site C:Day 3-Site A:Day 1  1.000000e-01 -1.16122806  1.36122806 1.0000000
Site D:Day 3-Site A:Day 1 -1.154632e-14 -1.26122806  1.26122806 1.0000000
Site E:Day 3-Site A:Day 1  2.000000e-01 -1.06122806  1.46122806 1.0000000
Site F:Day 3-Site A:Day 1 -8.666667e-01 -2.12789472  0.39456139 0.6055640
Site C:Day 1-Site B:Day 1  1.333333e-01 -1.12789472  1.39456139 1.0000000
Site D:Day 1-Site B:Day 1 -2.833333e-01 -1.54456139  0.97789472 0.9999983
Site E:Day 1-Site B:Day 1  4.666667e-01 -0.79456139  1.72789472 0.9982937
Site F:Day 1-Site B:Day 1 -5.333333e-01 -1.79456139  0.72789472 0.9919455
Site A:Day 2-Site B:Day 1  8.666667e-01 -0.39456139  2.12789472 0.6055640
Site B:Day 2-Site B:Day 1  3.483333e+00  2.22210528  4.74456139 0.0000000
Site C:Day 2-Site B:Day 1  5.000000e-01 -0.76122806  1.76122806 0.9961158
Site D:Day 2-Site B:Day 1  6.333333e-01 -0.62789472  1.89456139 0.9547996
Site E:Day 2-Site B:Day 1  1.183333e+00 -0.07789472  2.44456139 0.0962409
Site F:Day 2-Site B:Day 1  5.833333e-01 -0.67789472  1.84456139 0.9793521
Site A:Day 3-Site B:Day 1 -2.833333e+00 -4.09456139 -1.57210528 0.0000000
Site B:Day 3-Site B:Day 1 -1.166667e-01 -1.37789472  1.14456139 1.0000000
Site C:Day 3-Site B:Day 1 -7.500000e-01 -2.01122806  0.51122806 0.8275762
Site D:Day 3-Site B:Day 1 -8.500000e-01 -2.11122806  0.41122806 0.6408077
Site E:Day 3-Site B:Day 1 -6.500000e-01 -1.91122806  0.61122806 0.9430958
Site F:Day 3-Site B:Day 1 -1.716667e+00 -2.97789472 -0.45543861 0.0003146
Site D:Day 1-Site C:Day 1 -4.166667e-01 -1.67789472  0.84456139 0.9995943
Site E:Day 1-Site C:Day 1  3.333333e-01 -0.92789472  1.59456139 0.9999814
Site F:Day 1-Site C:Day 1 -6.666667e-01 -1.92789472  0.59456139 0.9293674
Site A:Day 2-Site C:Day 1  7.333333e-01 -0.52789472  1.99456139 0.8524089
Site B:Day 2-Site C:Day 1  3.350000e+00  2.08877194  4.61122806 0.0000000
Site C:Day 2-Site C:Day 1  3.666667e-01 -0.89456139  1.62789472 0.9999280
Site D:Day 2-Site C:Day 1  5.000000e-01 -0.76122806  1.76122806 0.9961158
Site E:Day 2-Site C:Day 1  1.050000e+00 -0.21122806  2.31122806 0.2477066
Site F:Day 2-Site C:Day 1  4.500000e-01 -0.81122806  1.71122806 0.9989121
Site A:Day 3-Site C:Day 1 -2.966667e+00 -4.22789472 -1.70543861 0.0000000
Site B:Day 3-Site C:Day 1 -2.500000e-01 -1.51122806  1.01122806 0.9999998
Site C:Day 3-Site C:Day 1 -8.833333e-01 -2.14456139  0.37789472 0.5699008
Site D:Day 3-Site C:Day 1 -9.833333e-01 -2.24456139  0.27789472 0.3629488
Site E:Day 3-Site C:Day 1 -7.833333e-01 -2.04456139  0.47789472 0.7717356
Site F:Day 3-Site C:Day 1 -1.850000e+00 -3.11122806 -0.58877194 0.0000507
Site E:Day 1-Site D:Day 1  7.500000e-01 -0.51122806  2.01122806 0.8275762
Site F:Day 1-Site D:Day 1 -2.500000e-01 -1.51122806  1.01122806 0.9999998
Site A:Day 2-Site D:Day 1  1.150000e+00 -0.11122806  2.41122806 0.1245536
Site B:Day 2-Site D:Day 1  3.766667e+00  2.50543861  5.02789472 0.0000000
Site C:Day 2-Site D:Day 1  7.833333e-01 -0.47789472  2.04456139 0.7717356
Site D:Day 2-Site D:Day 1  9.166667e-01 -0.34456139  2.17789472 0.4985047
Site E:Day 2-Site D:Day 1  1.466667e+00  0.20543861  2.72789472 0.0064733
Site F:Day 2-Site D:Day 1  8.666667e-01 -0.39456139  2.12789472 0.6055640
Site A:Day 3-Site D:Day 1 -2.550000e+00 -3.81122806 -1.28877194 0.0000000
Site B:Day 3-Site D:Day 1  1.666667e-01 -1.09456139  1.42789472 1.0000000
Site C:Day 3-Site D:Day 1 -4.666667e-01 -1.72789472  0.79456139 0.9982937
Site D:Day 3-Site D:Day 1 -5.666667e-01 -1.82789472  0.69456139 0.9846262
Site E:Day 3-Site D:Day 1 -3.666667e-01 -1.62789472  0.89456139 0.9999280
Site F:Day 3-Site D:Day 1 -1.433333e+00 -2.69456139 -0.17210528 0.0092848
Site F:Day 1-Site E:Day 1 -1.000000e+00 -2.26122806  0.26122806 0.3318621
Site A:Day 2-Site E:Day 1  4.000000e-01 -0.86122806  1.66122806 0.9997638
Site B:Day 2-Site E:Day 1  3.016667e+00  1.75543861  4.27789472 0.0000000
Site C:Day 2-Site E:Day 1  3.333333e-02 -1.22789472  1.29456139 1.0000000
Site D:Day 2-Site E:Day 1  1.666667e-01 -1.09456139  1.42789472 1.0000000
Site E:Day 2-Site E:Day 1  7.166667e-01 -0.54456139  1.97789472 0.8750337
Site F:Day 2-Site E:Day 1  1.166667e-01 -1.14456139  1.37789472 1.0000000
Site A:Day 3-Site E:Day 1 -3.300000e+00 -4.56122806 -2.03877194 0.0000000
Site B:Day 3-Site E:Day 1 -5.833333e-01 -1.84456139  0.67789472 0.9793521
Site C:Day 3-Site E:Day 1 -1.216667e+00 -2.47789472  0.04456139 0.0733605
Site D:Day 3-Site E:Day 1 -1.316667e+00 -2.57789472 -0.05543861 0.0300785
Site E:Day 3-Site E:Day 1 -1.116667e+000.95  0.14456139 0.1589473
Site F:Day 3-Site E:Day 1 -2.183333e+00 -3.44456139 -0.92210528 0.0000003
Site A:Day 2-Site F:Day 1  1.400000e+00  0.13877194  2.66122806 0.0131744
Site B:Day 2-Site F:Day 1  4.016667e+00  2.75543861  5.27789472 0.0000000
Site C:Day 2-Site F:Day 1  1.033333e+00 -0.22789472  2.29456139 0.2741465
Site D:Day 2-Site F:Day 1  1.166667e+00 -0.09456139  2.42789472 0.1096751
Site E:Day 2-Site F:Day 1  1.716667e+00  0.45543861  2.97789472 0.0003146
Site F:Day 2-Site F:Day 1  1.116667e+00 -0.14456139  2.37789472 0.1589473
Site A:Day 3-Site F:Day 1 -2.300000e+00 -3.56122806 -1.03877194 0.0000000
Site B:Day 3-Site F:Day 1  4.166667e-01 -0.84456139  1.67789472 0.9995943
Site C:Day 3-Site F:Day 1 -2.166667e-01 -1.47789472  1.04456139 1.0000000
Site D:Day 3-Site F:Day 1 -3.166667e-01 -1.57789472  0.94456139 0.9999912
Site E:Day 3-Site F:Day 1 -1.166667e-01 -1.37789472  1.14456139 1.0000000
Site F:Day 3-Site F:Day 1 -1.183333e+00 -2.44456139  0.07789472 0.0962409
Site B:Day 2-Site A:Day 2  2.616667e+00  1.35543861  3.87789472 0.0000000
Site C:Day 2-Site A:Day 2 -3.666667e-01 -1.62789472  0.89456139 0.9999280
Site D:Day 2-Site A:Day 2 -2.333333e-01 -1.49456139  1.02789472 0.9999999
Site E:Day 2-Site A:Day 2  3.166667e-01 -0.94456139  1.57789472 0.9999912
Site F:Day 2-Site A:Day 2 -2.833333e-01 -1.54456139  0.97789472 0.9999983
Site A:Day 3-Site A:Day 2 -3.700000e+00 -4.96122806 -2.43877194 0.0000000
Site B:Day 3-Site A:Day 2 -9.833333e-01 -2.24456139  0.27789472 0.3629488
Site C:Day 3-Site A:Day 2 -1.616667e+00 -2.87789472 -0.35543861 0.0011253
Site D:Day 3-Site A:Day 2 -1.716667e+00 -2.97789472 -0.45543861 0.0003146
Site E:Day 3-Site A:Day 2 -1.516667e+00 -2.77789472 -0.25543861 0.0036952
Site F:Day 3-Site A:Day 2 -2.583333e+00 -3.84456139 -1.32210528 0.0000000
Site C:Day 2-Site B:Day 2 -2.983333e+00 -4.24456139 -1.72210528 0.0000000
Site D:Day 2-Site B:Day 2 -2.850000e+00 -4.11122806 -1.58877194 0.0000000
Site E:Day 2-Site B:Day 2 -2.300000e+00 -3.56122806 -1.03877194 0.0000000
Site F:Day 2-Site B:Day 2 -2.900000e+00 -4.16122806 -1.63877194 0.0000000
Site A:Day 3-Site B:Day 2 -6.316667e+00 -7.57789472 -5.05543861 0.0000000
Site B:Day 3-Site B:Day 2 -3.600000e+00 -4.86122806 -2.33877194 0.0000000
Site C:Day 3-Site B:Day 2 -4.233333e+00 -5.49456139 -2.97210528 0.0000000
Site D:Day 3-Site B:Day 2 -4.333333e+00 -5.59456139 -3.07210528 0.0000000
Site E:Day 3-Site B:Day 2 -4.133333e+00 -5.39456139 -2.87210528 0.0000000
Site F:Day 3-Site B:Day 2 -5.200000e+00 -6.46122806 -3.93877194 0.0000000
Site D:Day 2-Site C:Day 2  1.333333e-01 -1.12789472  1.39456139 1.0000000
Site E:Day 2-Site C:Day 2  6.833333e-01 -0.57789472  1.94456139 0.9134954
Site F:Day 2-Site C:Day 2  8.333333e-02 -1.17789472  1.34456139 1.0000000
Site A:Day 3-Site C:Day 2 -3.333333e+00 -4.59456139 -2.07210528 0.0000000
Site B:Day 3-Site C:Day 2 -6.166667e-01 -1.87789472  0.64456139 0.9646272
Site C:Day 3-Site C:Day 2 -1.250000e+00 -2.51122806  0.01122806 0.0551896
Site D:Day 3-Site C:Day 2 -1.350000e+00 -2.61122806 -0.08877194 0.0218071
Site E:Day 3-Site C:Day 2 -1.150000e+00 -2.41122806  0.11122806 0.1245536
Site F:Day 3-Site C:Day 2 -2.216667e+00 -3.47789472 -0.95543861 0.0000002
Site E:Day 2-Site D:Day 2  5.500000e-01 -0.71122806  1.81122806 0.9887628
Site F:Day 2-Site D:Day 2 -5.000000e-02 -1.31122806  1.21122806 1.0000000
Site A:Day 3-Site D:Day 2 -3.466667e+00 -4.72789472 -2.20543861 0.0000000
Site B:Day 3-Site D:Day 2 -7.500000e-01 -2.01122806  0.51122806 0.8275762
Site C:Day 3-Site D:Day 2 -1.383333e+00 -2.64456139 -0.12210528 0.0156284
Site D:Day 3-Site D:Day 2 -1.483333e+00 -2.74456139 -0.22210528 0.0053838
Site E:Day 3-Site D:Day 2 -1.283333e+00 -2.54456139 -0.02210528 0.0409949
Site F:Day 3-Site D:Day 2 -2.350000e+00 -3.61122806 -1.08877194 0.0000000
Site F:Day 2-Site E:Day 2 -6.000000e-01 -1.86122806  0.66122806 0.9727496
Site A:Day 3-Site E:Day 2 -4.016667e+00 -5.27789472 -2.75543861 0.0000000
Site B:Day 3-Site E:Day 2 -1.300000e+00 -2.56122806 -0.03877194 0.0351682
Site C:Day 3-Site E:Day 2 -1.933333e+00 -3.19456139 -0.67210528 0.0000151
Site D:Day 3-Site E:Day 2 -2.033333e+00 -3.29456139 -0.77210528 0.0000033
Site E:Day 3-Site E:Day 2 -1.833333e+00 -3.09456139 -0.57210528 0.0000642
Site F:Day 3-Site E:Day 2 -2.900000e+00 -4.16122806 -1.63877194 0.0000000
Site A:Day 3-Site F:Day 2 -3.416667e+00 -4.67789472 -2.15543861 0.0000000
Site B:Day 3-Site F:Day 2 -7.000000e-01 -1.96122806  0.56122806 0.8953977
Site C:Day 3-Site F:Day 2 -1.333333e+00 -2.59456139 -0.07210528 0.0256487
Site D:Day 3-Site F:Day 2 -1.433333e+00 -2.69456139 -0.17210528 0.0092848
Site E:Day 3-Site F:Day 2 -1.233333e+00 -2.49456139  0.02789472 0.0637326
Site F:Day 3-Site F:Day 2 -2.300000e+00 -3.56122806 -1.03877194 0.0000000
Site B:Day 3-Site A:Day 3  2.716667e+00  1.45543861  3.97789472 0.0000000
Site C:Day 3-Site A:Day 3  2.083333e+00  0.82210528  3.34456139 0.0000015
Site D:Day 3-Site A:Day 3  1.983333e+00  0.72210528  3.24456139 0.0000071
Site E:Day 3-Site A:Day 3  2.183333e+00  0.92210528  3.44456139 0.0000003
Site F:Day 3-Site A:Day 3  1.116667e+00 -0.14456139  2.37789472 0.1589473
Site C:Day 3-Site B:Day 3 -6.333333e-01 -1.89456139  0.62789472 0.9547996
Site D:Day 3-Site B:Day 3 -7.333333e-01 -1.99456139  0.52789472 0.8524089
Site E:Day 3-Site B:Day 3 -5.333333e-01 -1.79456139  0.72789472 0.9919455
Site F:Day 3-Site B:Day 3 -1.600000e+00 -2.86122806 -0.33877194 0.0013803
Site D:Day 3-Site C:Day 3 -1.000000e-01 -1.36122806  1.16122806 1.0000000
Site E:Day 3-Site C:Day 3  1.000000e-01 -1.16122806  1.36122806 1.0000000
Site F:Day 3-Site C:Day 3 -9.666667e-01 -2.22789472  0.29456139 0.3953438
Site E:Day 3-Site D:Day 3  2.000000e-01 -1.06122806  1.46122806 1.0000000
Site F:Day 3-Site D:Day 3 -8.666667e-01 -2.12789472  0.39456139 0.6055640
Site F:Day 3-Site E:Day 3 -1.066667e+00 -2.32789472  0.19456139 0.2229553
```
