## Supporting Information Appendix S4 for "Simultaneous passive acoustic monitoring uncovers evidence of potentially overlooked temporal variation in an Amazonian bird community"

**Supporting Information Appendix S4:** Model covariate summary statistics, including smoothing parameters (*k*), effective degrees of freedom (*edf*), *F*-statistics, and *p*-values. Covariate effects highlighted in bold indicate statistically significant differences (*p* < 0.05) among groups based on analysis of variance (ANOVA).

|  | **Model summary** | | | **ANOVA** | |
| --- | --- | --- | --- | --- | --- |
| **Covariate** | **Estimate ± SE** | ***k*** | ***edf*** | ***F*** | ***p*-value** |
| Species richness model (*R^2^* = 0.637) | | | | | |
| **Intercept** | 1.44 ± 0.05 |  |  |  | **< 0.001** |
| s(Site, by = Day) |  |  |  |  |  |
| Day 1 |  | 6.00 | 0.00 | 0.00 | 0.952 |
| **Day 2** |  | 6.00 | 4.78 | 4.52 | **< 0.001** |
| **Day 3** |  | 6.00 | 4.67 | 3.12 | **0.001** |
| s(Minute, by = Recording) |  |  |  |  |  |
| A1 |  | 9.00 | 1.00 | 2.16 | 0.142 |
| A2 |  | 9.00 | 1.00 | 0.70 | 0.402 |
| A3 |  | 9.00 | 1.00 | 1.38 | 0.239 |
| **B1** |  | 9.00 | 2.22 | 6.31 | **0.001** |
| **B2** |  | 9.00 | 1.00 | 4.743 | **0.029** |
| **B3** |  | 9.00 | 2.27 | 10.60 | **< 0.001** |
| **C1** |  | 9.00 | 1.00 | 18.11 | **< 0.001** |
| C2 |  | 9.00 | 1.93 | 1.365 | 0.295 |
| **C3** |  | 9.00 | 1.70 | 11.29 | **0.002** |
| D1 |  | 9.00 | 1.73 | 0.81 | 0.312 |
| D2 |  | 9.00 | 1.00 | 0.02 | 0.894 |
| **D3** |  | 9.00 | 2.34 | 5.24 | **0.002** |
| **E1** |  | 9.00 | 1.73 | 14.73 | **< 0.001** |
| **E2** |  | 9.00 | 1.00 | 4.05 | **0.045** |
| **E3** |  | 9.00 | 1.00 | 14.61 | **< 0.001** |
| **F1** |  | 9.00 | 2.23 | 6.87 | **< 0.001** |
| F2 |  | 9.00 | 2.27 | 2.13 | 0.251 |
| **F3** |  | 9.00 | 1.00 | 11.62 | **< 0.001** |
| Total vocal prevalence model (*R^2^* = 0.829) | | | | | |
| **Intercept** | 2.91 ± 0.04 |  |  |  | **< 0.001** |
| s(Site, by = Day) |  |  |  |  |  |
| **Day 1** |  | 6.00 | 3.65 | 2.43 | **0.001** |
| **Day 2** |  | 6.00 | 5.88 | 88.78 | **< 0.001** |
| **Day 3** |  | 6.00 | 5.84 | 33.68 | **< 0.001** |
| s(Minute, by = Recording) |  |  |  |  |  |
| **A1** |  | 9.00 | 7.41 | 9.88 | **< 0.001** |
| **A2** |  | 9.00 | 2.36 | 3.99 | **0.009** |
| **A3** |  | 9.00 | 7.78 | 10.33 | **< 0.001** |
| **B1** |  | 9.00 | 5.63 | 10.76 | **< 0.001** |
| **B2** |  | 9.00 | 1.00 | 23.74 | **< 0.001** |
| **B3** |  | 9.00 | 4.55 | 26.92 | **< 0.001** |
| **C1** |  | 9.00 | 2.68 | 56.14 | **< 0.001** |
| **C2** |  | 9.00 | 4.60 | 19.22 | **< 0.001** |
| **C3** |  | 9.00 | 5.01 | 18.51 | **< 0.001** |
| **D1** |  | 9.00 | 5.46 | 11.31 | **< 0.001** |
| **D2** |  | 9.00 | 7.18 | 10.54 | **< 0.001** |
| **D3** |  | 9.00 | 5.02 | 12.36 | **< 0.001** |
| **E1** |  | 9.00 | 6.56 | 28.49 | **< 0.001** |
| **E2** |  | 9.00 | 4.78 | 16.41 | **< 0.001** |
| **E3** |  | 9.00 | 1.00 | 164.49 | **< 0.001** |
| **F1** |  | 9.00 | 3.80 | 43.03 | **< 0.001** |
| **F2** |  | 9.00 | 6.42 | 11.92 | **< 0.001** |
| **F3** |  | 9.00 | 3.09 | 27.97 | **< 0.001** |
